# Structural disruption of genomic regions containing ultraconserved elements is associated with neurodevelopmental phenotypes

**DOI:** 10.1101/233197

**Authors:** Ruth B. McCole, Wren Saylor, Claire Redin, Chamith Y. Fonseka, Harrison Brand, Jelena Erceg, Michael E. Talkowski, C.-ting Wu

## Abstract

The development of the human brain and nervous system can be affected by genetic or environmental factors. Here we focus on characterizing the genetic perturbations that accompany and may contribute to neurodevelopmental phenotypes. Specifically, we examine two types of structural variants, namely, copy number variation and balanced chromosome rearrangements, discovered in subjects with neurodevelopmental disorders and related phenotypes. We find that a feature uniting these types of genetic aberrations is a proximity to ultraconserved elements (UCEs), which are sequences that are perfectly conserved between the reference genomes of distantly related species. In particular, while UCEs are generally depleted from copy number variant regions in healthy individuals, they are, on the whole, enriched in genomic regions disrupted by copy number variants or breakpoints of balanced rearrangements in affected individuals. Additionally, while genes associated with neurodevelopmental disorders are enriched in UCEs, this does not account for the excess of UCEs either in copy number variants or close to the breakpoints of balanced rearrangements in affected individuals. Indeed, our data are consistent with some manifestations of neurodevelopmental disorders resulting from a disruption of genome integrity in the vicinity of UCEs.

## Introduction

The etiology of neurodevelopmental disorders (NDDs) involves both genetic and environmental factors. In this study, we are concerned with the genetic, rather than environmental, disruptions that may result in NDDs including autism spectrum disorder (ASD) and other neurodevelopmental phenotypes. Evidence for a large genetic component to ASD etiology comes from studies over many decades showing a much higher concordance for ASD in monozygotic twins when compared with dizygotic twins (reviewed in Huguet *et al.*^1^). More generally, NDDs have been associated with an increasingly large range of genetic disruptions that are highly diverse in genomic location, type, and origin and include single nucleotide changes, copy number variations (CNVs), and complex, sometimes copy number neutral, chromosomal rearrangements (reviewed in Kloosterman *et al.*^2^, Hu *et al.*^3^, and Sahin *et al.*^4^). With regard to the origin of NDD-associated genomic aberrations, *de novo* events have recently been a major focus, although somatic events during development and inherited variation have also been examined (reviewed in Ronemus *et al.*^5^ and Hu *et al.*^3^).

Since genetic alterations found in subjects with NDDs and related phenotypes vary in genomic position, aberration type, and origin, we wondered whether examining these aberrations from the perspective of comparative genomics might unveil commonalities among this set of genomic abnormalities. We chose this approach in light of the fact that high sequence conservation is a hallmark of neurodevelopmental^6^, presynaptic, and synaptic genes^7, 8^. Specifically, this study queries the relationship between genomic rearrangements in individuals with NDDs or related phenotypes and a set of human sequences known as ultraconserved elements (UCEs). Examples of highly conserved genes with relevance to neurodevelopment and that also contain UCEs include AUTS2^9^, FOXP2^10^, and BCL11A^11^.

First reported in 2004, human UCEs are defined as regions of the human genome displaying extreme sequence conservation with distantly related vertebrates ^12–14^. They comprise one of the most enigmatic aspects of our genomes, as the underlying reasons for ultraconservation are still being debated; while UCEs contribute to gene regulation, contain many transcription factor binding motifs, are often transcribed, and may produce phenotypes when mutated or deleted, these properties are not universally considered sufficient to explain ultraconservation^14–30^, reviewed in Elgar *et al.* ^31^ and Harmston *et al.*^32^. We suggest that ultraconservation may derive from a mechanism in which maternal and paternal copies of UCEs are compared within the nucleus, such that individuals bearing discrepancies in sequence or copy number of UCEs or rearrangements that disrupt the comparison have reduced fitness^33–35^(see also Elgar *et al.*^31^ and Kritsas *et al.*^36^). Such a process has the potential of reducing the burden in germline, embryonic, and adult somatic cell lineages of cells carrying deleterious genomic aberrations. Interestingly, the positioning and clustering of UCEs along chromosomes can be highly conserved^13, 37–41^, and regions of the genome that include highly conserved elements appear to interact in three-dimensional space^42^. These observations are in line with the possibility that UCEs have a function that requires their specific three-dimensional positioning within the nucleus, such as the pairing of homologous genomic regions containing UCEs. Importantly, ample evidence exists for the capacity of vertebrate cells to support homolog pairing outside as well as during meiosis (reviewed in Joyce *et al.*^43^ and references within) and a role for pairing in gene regulation, sequence comparison, and copy counting (e.g. Joyce *et al.*^43^ Hammond *et al.*^44^, Gladyshev *et al.*^45^, and references within). Our model is consistent with the viability and fertility observed by Ahituv *et al.*^46^ of mice lacking both copies of any of four noncoding UCEs, as deletion of both copies of a UCE would preclude detection of these deletions. It is also compatible with additional, gene regulatory functions for UCEs and therefore is not contradicted by discoveries of abnormal phenotypes, such as improper eye development^25^, that arise from mutations in UCEs.

This model predicts that changes to UCE copy number would be disfavored in healthy cells, and consistent with this, meta-analyses of over two dozen datasets of copy number variations (CNVs) representing healthy cells showed a strong depletion of UCEs from deleted or duplicated regions^33–35, 47^, while a study of plants has shown that sequences conserved in distantly related plant genomes are depleted from segmental duplications^36^. The model also predicts that disruption of UCE copy number above the low levels seen in healthy tissues would be associated with disease, and we have found evidence for this from our meta-analysis of seventeen datasets of CNVs derived from cancer tissues. In particular, we observed a higher level of UCE disruption by cancer-specific CNVs than by CNVs in healthy tissues, with a number of cancer CNV datasets showing an enrichment for UCEs^35^. Of special relevance to NDD, Martinez *et al.*,^48^ pursued our prediction that abnormal dosage of UCEs would be associated with lowered fitness^33–35^ by querying whether the relationship between CNVs and UCEs in individuals with mental delay and congenital abnormalities would differ from that in healthy individuals. They showed that there was indeed an association between UCE position and the structural variants discovered in the genomes of these patients.

Our previous studies all focused on elucidating the positional relationship between UCEs and CNVs, but did not address the relationship between UCEs and copy number neutral rearrangements. The present study also examines CNVs and then, for the first time, addresses copy number neutral rearrangements, taking advantage of the large number of new datasets describing structural variants in subjects with NDD or related phenotypes. Thus, in addition to elucidating the positional relationship between UCEs and rearrangement breakpoints, this study tackles the prediction that genomic disruptions that compromise the comparison process of UCEs will be associated with disease. We begin by analyzing CNVs in a first cohort with respect to their origin, separating inherited CNVs from *de novo* CNVs that are, as a group, associated with NDD causation^5, 49, 50^. We then compare *de novo* CNVs in a separate ASD cohort to *de novo* CNVs discovered in unaffected siblings. Finally, we examine inversion, translocation, and complex rearrangements in subjects with NDDs or related phenotypes.

Our results demonstrate a striking departure from the depletion of UCEs observed in regions affected by CNVs in healthy individuals. Indeed, individuals with NDD or related phenotypes reveal an enrichment of UCEs in genomic regions encompassed by CNVs or flanking the breakpoints of balanced rearrangements. Importantly, while UCEs are also prevalent within genes whose disruption is thought to elevate NDD risk, this prevalence is not sufficient to explain the excess of UCEs encompassed in the CNVs, nor the extent of the enrichment of UCEs near the breakpoints of balanced rearrangements, in affected cohorts. In summary, our findings suggest three non-conflicting possibilities: 1) UCEs may signpost new candidate genes and critical regions for NDD involvement, as has been suggested previously^48^; 2) disruption of either UCE dosage or nuclear position may itself be a causal factor in NDD etiology; and 3) UCEs may function to support genomic integrity, with NDD being one possible outcome when they are compromised.

## Materials and Methods

### Datasets

All datasets are available from the publications or web resources listed and *per* these sources the procedures followed were in accordance with the ethical standards of the responsible committee on human experimentation and proper informed consent was obtained.

#### Vulto-van Silfhout ASD *de novo* CNVs and Vulto-van Silfhout ASD inherited CNVs

(See Figure S1): CNV regions corresponding to subjects with autism spectrum disorder were taken from Vulto van-Silfhout *et al.*^51^. Out of 1663 events, 206 events corresponded to subjects with ASD phenotype. Of these, 54 were *de novo*, and 94 were inherited. When overlapping events were combined, 45 regions made up our dataset ‘Vulto-van Silfhout ASD *de novo* CNVs’, and 81 regions comprised our dataset ‘Vulto-van Silfhout ASD inherited CNVs’.

#### Sanders ASD *de novo* CNVs and Sanders sibling *de novo* CNVs (See Figure S1)

CNVs were drawn from Sanders *et al.* ^52^. Inherited CNVs were not considered, because only rare inherited CNVs were available from Sanders *et al.* and considering only a subset of all inherited CNVs would not provide us with a full representation of their relationships to UCEs. For *de novo* CNVs, there were 495 events for cases and 121 events for controls. These were filtered to retain validated events, leaving 328 case events and 90 control events. Whole chromosome aneuploidies, as well as one event covering >94% of the Y chromosome q arm, were removed. For cases, 321 events remained, which when overlapping events were combined, produced 191 regions, which we call ‘Sanders ASD *de novo* CNVs’. For controls, 86 events remained, which, when overlapping events were combined, produced 79 regions that we refer to as ‘Sanders sibling *de novo* CNVs’ (See Figure S1).

#### NDD breakpoints set 1 (See Figure S3)

Breakpoints were taken from 5 publications: Granot-Hershkovitz *et al.* 2011 ^53^, Kloosterman *et al.* 2011 ^54^, Chiang *et al.* 2012 ^55^, Talkowski *et al.* 2012 ^56^, and Nazaryan *et al.* 2014 ^57^ (see Table S1 and Figure S3). Only breakpoints documented to accompany ≤1kb of inserted or deleted DNA were included; we did not place any limits on the amount of DNA rearranged by the variant. Duplicate breakpoints and breakpoints from subjects without neurodevelopmental phenotypes were removed. Where necessary, coordinates were converted to genome build hg18 using the UCSC liftover tool (http://genome.ucsc.edu/cgi-bin/hgLiftOver). The final ‘NDD breakpoints set 1’ dataset comprised 157 breakpoints. Flanking regions of 100kb on either side of each breakpoint were added using the Bedtools ^58^ slop tool, implemented using Pybedtools ^59^, producing dataset ‘NDD breakpoints set 1 with 100kb flanks’, consisting of 76 regions.

#### NDD breakpoints Set 2 (See Figure S3)

Breakpoints were taken from Redin *et al.*^60^. 1858 breakpoints were filtered to retain only those where the total genomic imbalance per individual is ≤1kb; we did not place any limits on the amount of DNA rearranged by the variant. These breakpoints were filtered to retain only those where the breakpoint was confirmed by capillary sequencing, resulting in 744 breakpoints. To produce a set of breakpoints that are different from those in NDD breakpoints Set 1, we removed any breakpoints listed by Redin *et al.*^60^ that were included in NDD breakpoints Set 1, leaving 590 breakpoints. We then filtered to obtain only breakpoints from patients reported as having an ear, eye, or nervous system phenotype, leaving 478 breakpoints. We also filtered to retain breakpoints from subjects with neither ear, nor eye, nor nervous system phenotypes, and these breakpoints formed our ‘Non-NDD breakpoint’ dataset, described below. Duplicate breakpoints were removed, leaving 453 breakpoints, which comprised our final dataset ‘NDD breakpoints set 2’. Flanking regions of 100kb on either side of each breakpoint were added using the Bedtools ^58^ slop tool, implemented using Pybedtools ^59^, and overlapping regions were combined to produce 224 regions, which we refer to as ‘NDD breakpoints Set 2 with 100kb flanks’.

#### Pooled NDD breakpoints (See Figure S3)

We combined the 157 breakpoints in NDD breakpoints set 1 with the 453 breakpoints in NDD breakpoint set 2 to create our dataset Pooled NDD breakpoints’ of 610 breakpoints. Flanking regions of 20kb, 50kb, 100kb, 500kb, and 1Mb on either side of each breakpoint were added using the Bedtools^58^ slop tool, implemented using Pybedtools^59^, and overlapping regions were combined to produce five new datasets, ‘Pooled NDD breakpoints with 20kb flanks’ containing 316 regions, ‘Pooled NDD breakpoints with 50kb flanks’ with 310 regions, ‘Pooled NDD breakpoints with 100kb flanks’ with 296 regions, ‘Pooled NDD breakpoints with 500kb flanks’ with 250 regions, and ‘Pooled NDD breakpoints with 1Mb flanks’, with 219 regions. To examine the distances between UCEs and Pooled NDD breakpoints, we allocated any breakpoint in our Pooled NDD breakpoint set into a cluster if it occurred within 1kb of any other breakpoint. We then retained just one breakpoint from each cluster, producing our dataset ‘Pooled NDD breakpoints within 1kb clustered’, which contained 331 breakpoints.

#### Pathogenic, likely pathogenic, and VUS breakpoints (See Figure S3)

We divided our NDD breakpoints set 2 according to the category of pathogenicity the breakpoints for each patient fell into. The categories were provided in Redin *et al.*^60^, and consisted of Pathogenic, Likely Pathogenic, and Variants of Unknown Significance (VUS). In total, 64 were from patients classified as having pathogenic variants, 150 were classified as having likely pathogenic variants, and 264 were from patients with variants of unknown significance. When 500kb flanks were added to either side of each breakpoint and overlapping regions were combined, the breakpoints made up the datasets ‘Pathogenic breakpoints with 500kb flanks’ with 32 regions, ‘Likely pathogenic breakpoints with 500kb flanks’ with 62 regions, and ‘VUS breakpoints with 500kb flanks’ with 120 regions.

#### Non-NDD breakpoints (See Figure S3)

Breakpoints from Redin *et al.*^60^ were filtered as described above and by phenotype to retain 76 breakpoints from subjects with neither ear, nor eye, nor nervous system phenotypes. 500kb flanks were added to these breakpoints to produce 38 regions.

#### Loss-of-function (LoF) genes

Genes with two or more loss-of-function mutations in ASD patients were collated by Redin *et al.* ^60^. The coordinates were converted from build hg19 to build hg18 using the UCSC liftover tool (http://genome.ucsc.edu/cgi-bin/hgLiftOver). The final gene list comprised 304 genes and is referred to as ‘LoF genes’ (see Table S1).

#### Constrained genes

We obtained all gene identifiers from Lek *et al.* ^61^ that had pLI score > 0.9, indicating these genes are loss-of-function intolerant in humans and likely to be selectively constrained. These identifiers were used to obtain coordinates for each gene in human genome build hg19, using the UCSC table browser (http://genome.ucsc.edu/cgi-bin/hgTables) and track ‘GENCODE genes V 19’. The coordinates were then lifted over to build hg18 using the UCSC liftover tool (http://genome.ucsc.edu/cgi-bin/hgLiftOver). The final list comprised 3249 genes and is called ‘Constrained genes’ (See Table S1).

#### Embryonic brain genes

We collected genes that showed higher expression in the embryonic human brain when compared to the postnatal human brain, identified by Iossifov *et al.*^62^. The coordinates were lifted over to build hg18 using the UCSC liftover tool (http://genome.ucsc.edu/cgi-bin/hgLiftOver). The final dataset, named ‘ Embryonic brain genes’ comprised 1903 genes (See Table S1).

#### Type 2 diabetes (T2D) genes

We obtained genes associated with type 2 diabetes from Morris *et al.*^63^, matching the gene symbols provided with gene coordinates in build hg18 using the UCSC table browser (http://genome.ucsc.edu/cgi-bin/hgTables) and manual searches on the UCSC genome browser (http://genome.ucsc.edu). In total, our ‘T2D genes’ dataset is made up of 70 genes (See Table S1).

#### Schizophrenia (SZ) genes

Genes implicated in schizophrenia risk by means of genome wide association studies (GWAS) were collected from the NHGRI-EBI GWAS catalog^64^ at http://www.ebi.ac.uk/gwas/search?query=Schizophrenia. Genes were filtered to retain those labeled ‘schizophrenia’; genes with label ‘schizophrenia and autism’ were not included.

#### Enhancers

Enhancers dataset was from the ENCODE combined genome segmentation from the ENCODE UCSC hub^65^ ‘E’ (enhancer) class genomic regions for six ENCODE cell/tissue types.

#### Human accelerated regions (HARs)

Genome coordinates for HARs were obtained from Doan *et al.*^66^, and lifted over to hg18 using the UCSC liftover tool (http://genome.ucsc.edu/cgi-bin/hgLiftOver).

#### Repetitive elements

Genomic coordinates for repetitive elements were obtained from the UCSC genome browser Repeat Masker track.

#### UCEs

Our UCE dataset comprises 896 regions that display 100% DNA conservation between all members of these three groups: 1) human, mouse, and rat; 2) human, dog, and mouse, and 3) human and chicken. These sequences are available in McCole *et al.*^35^ and are provided in Table S1. We divide the UCEs into intergenic, intronic, and exonic elements, defining introns and exons using the UCSC known genes track for genome build hg18. Exonic elements are those UCEs that wholly or partially overlap exons, and comprise 179 elements. Intronic elements are any of the non-exonic UCEs that wholly or partially overlap introns, and comprise 416 elements. The remaining 301 UCEs are categorized as intergenic.

### Depletion and enrichment analysis

To determine depletion or enrichment of UCEs in genomic regions of interest, we use a similar method as previously described^33–35^. We begin by merging all overlapping genomic regions of interest, using a custom python script, which functions almost identically to the BEDtools^58^ merge function. We then remove from these merged genomic regions of interest any overlaps with unsequenced portions of the human genome. Next we calculate the overlap in bp between merged genomic regions of interest and each of the three categories of UCEs (intergenic, intronic, exonic). For example, for the intergenic UCEs, the pipeline first calculates the overlap in basepairs between the intergenic UCEs and CNVs (or other regions of interest), which we refer to as the ‘observed overlap’. Next, the pipeline randomly permutes the positions of the UCEs within the appropriate portion of the genome; with, for example, the intergenic UCEs being permuted only within intergenic regions. The pipeline then calculates the overlap between the CNVs and the randomly positioned elements that are matched in number, length, and genomic region, and we call this the ‘expected overlap’. The randomization process is then repeated 1,000 times to generate a distribution of expected overlaps.

When analyzing the entire UCE set, this process is repeated for intronic and exonic UCEs and then the intergenic, intronic, and exonic observed overlaps are added to produce the observed overlap for the entire UCE set, and similarly, expected overlaps for intergenic, intronic, and exonic regions within each of the 1,000 permutations are added to produce a distribution of expected overlaps for the entire UCE set. The ratio of observed/expected overlap is reported as ‘obs/exp’. To assess statistical significance, the distribution of expected overlaps is assessed for normality using a Kolomogorov-Smirnov (KS) test, and the resulting P-value is reported. If the distribution is consistent with normality, a Z-test is used to determine the statistical significance of the depletion (obs/exp < 1) or enrichment (obs/exp > 1) of observed UCE overlap with genomic regions of interest, compared with expected overlap, and the resulting P value is reported. P values below 10^-17^ are not calculated exactly by the test and so are reported as here as <1×10^−17^. Occasionally, our distribution of expected overlaps is not normally distributed, precluding the generation of a P-value and, in this situation, we provide instead the proportion of expected overlaps that equaled or exceeded the observed overlap. To allow for the possibility of either enrichment or depletion, a two-tailed test is employed with a combined α of 0.05.

### Binomial test for overlap of Pooled NDD breakpoints with UCEs

To calculate the probability of observing 1 or more breakpoints overlapping UCEs, we calculated that UCEs cover 241,287bp. Assuming a 3 Gb human genome, the proportion of the human genome that UCEs make up is 0.00008. We then performed a binomial test with the probability of success set to 0.00008 and the number of trials equal to the number of breakpoints.

### Distances from breakpoints to UCEs

UCEs within 1kb of each other were grouped and, for each group, one member of the group was chosen randomly for analysis, to avoid biasing the results with very closely clustered UCEs. Distances between UCEs and the nearest breakpoint were calculated using a custom python script utilizing Pybedtools^59^. The resulting distances were then expressed as frequencies in the form of a histogram. Additionally, 1,000 sets of regions matched to UCEs in terms of length, number, and genomic portion (intergenic, intronic, or exonic), which we call ‘control regions’ were generated using our depletion/enrichment analysis pipeline. The distances between these control regions and their nearest breakpoint was then calculated. The distribution of UCE-breakpoint distances, and of control-breakpoint distances, was compared using an Anderson-Darling test implemented in python using the scipy function anderson_ksamp, which provides exact significance thresholds and an approximated P value (stated). The maximum distance allowed was set at 5.2Mb, to match the distribution displayed in Figure 2E. For visualization of the results and comparison to the histogram of UCE-breakpoint distances, the distances in each of the 1,000 trials were expressed as frequencies and the interquartile range between the 1,000 trials at each distance was plotted.

### Depletion and enrichment analysis with gene sets excluded

We carried out this analysis using the same pipeline as for ‘Depletion and enrichment analysis’, above. We excluded the regions of the genome corresponding to our gene set of interest, as well as in some experiments the regions 100kb upstream and downstream of these genes. This means that any UCEs and regions of interest such as CNVs that fall within the excluded regions were not included in the analysis. Additionally, no UCE-matched random control regions were allowed to fall within the excluded regions.

### Partial correlation analysis

Data for enhancers, HARs, and repetitive elements was obtained as described above. GC percentage was calculated using the hg18 human reference genome and bedtools^58, 59^ ‘nuc’ function. Custom python scripts were used to calculate the density of genomic regions of interest in bp within bins of 100kb by summing the number of bp covered by the regions and dividing this by the genomic coverage of the bin (100kb). Bins containing unsequenced regions of the reference genome were excluded from the analysis. The Spearman partial correlation coefficient and accompanying P-value, which represents the partial correlation between UCE density and a feature of interest such as Pooled NDD breakpoints after removal of any contribution of any correlation with a third feature, such as enhancers, was then calculated using matlab function *partialcorr* implemented within custom python scripts.

### UCEs overlapping regions of interest (Table S3)

For each UCE, its ID, coordinates, type (intergenic, intronic, exonic), and the UCSC and HGNC gene symbol for any overlapping gene is listed. Then, for each dataset listed in Table S1, if the UCE is overlapped by this dataset, a ‘1’ is listed next to this UCE. If the UCE does not overlap this dataset, a ‘0’ is listed.

### Scripts

Custom python scripts associated with this study can be accessed at https://github.com/rmccole/UCEs_and_neurodevelopment

## Results

### *De novo* CNVs in cases of autism spectrum disorder disrupt UCE dosage

Our first analysis compared how *de novo* and inherited CNVs in the genomes of (ASD) probands are positioned relative to UCEs, noting that the bulk of the CNV-mediated genetic etiology of ASD is thought to be borne by *de novo*, rather than inherited, CNVs^5, 49, 50, 52, 67–72^. *De novo* CNVs are defined as being present in the genomes of the proband but not either parent, while inherited CNVs are defined as identifiable in the genomes of the proband and at least one parent. From Vulto-van Silfhout *et al.*^51^, we obtained 206 CNVs from patients with an ASD phenotype, of which 54 arose *de novo* and 94 were inherited; 58 were classed as “inheritance unknown” and were not examined further (see Figure S1 for details of filtering steps). When segments of overlapping *de novo* CNVs were combined, a final dataset of 45 distinct ‘Vulto-van Silfhout ASD *de novo* CNVs’ covering 5.91% of the genome was obtained (Table S1). Similarly, by merging overlapping inherited CNVs, we obtained 81 CNV regions covering 3.19% of the genome (Table S1), and we call these CNVs ‘Vulto-van Silfhout ASD inherited CNVs’.

We queried our datasets of CNVs for their relationship to the positions of UCEs using an analysis pipeline that builds upon our previously published methods^33–35^ (Methods, “Depletion and enrichment analysis”). The UCEs are first partitioned into intergenic, intronic, and exonic subsets of 301, 416, and 179 UCEs respectively^35^, and then the pipeline operates on the three categories of UCEs separately. Briefly, we calculate a distribution of expected overlaps that reflects the overlaps that would arise between the CNVs and UCEs if the UCEs were distributed randomly within the appropriate genomic subset (exonic UCEs being randomly placed within exons, for example). The observed overlap between UCEs and CNVs is then compared to the distribution of expected overlaps using a Z-test, with the outcome accompanied by a P-value and ratio of observed/expected (obs/exp) overlaps. Occasionally, our distribution of expected overlaps is not normally distributed, in which case we provide the proportion of expected overlaps that equaled or exceeded the observed overlap.

Because we generate a bespoke expected overlap distribution for each CNV dataset of interest, our analysis is suitable for analyzing and comparing datasets that differ widely with respect to their genomic coverage, as is common for datasets of CNVs and other structural variants. Additionally, our expected overlap distribution reflects the division of UCEs between the intergenic, intronic, and exonic regions of the genome, ensuring that any interesting differences between the observed and expected overlaps cannot merely be due to overall trends in UCE occupancy of different genomic regions (for example, 20% of the UCEs are exonic even though only approximately 3% of the human genome is composed of exons^73^). As for our dataset of UCEs, we made use of a previously defined set of 896 UCEs^33–35^ that are ≥200bp in length and 100% identical in DNA sequence between the reference genomes of human, mouse, and rat, or of human, dog, and mouse, or of human and chicken (Table S1) all of which represent unique sequences in the genome^33^.

We began our study with the expectation that overall depletion of UCEs from a set of CNVs is an indicator that such a set of CNVs is likely benign^33–35^. For example, our most recent analysis of healthy individuals, including 8 datasets of predominantly inherited CNVs covering 51.4% of the genome and 4 datasets of *de novo* CNVs covering 0.9% of the genome yielded obs/exp ratios of 0.771 (P = 1.7 × 10^−12^) and 0.395 (P = 0.044), respectively^35^.

Here, we used a 2-tailed α of 0.025 and found that the Vulto-van Silfhout ASD CNVs inherited from unaffected parents show statistically significant depletion (Figure 1A, Table S2A; P = 0.003, obs/exp = 0.493), with only 17 UCEs overlapping CNVs. This CNV profile is similar to our findings for CNVs representing healthy individuals^35^ (). In stark contrast, Vulto-van Silfhout ASD *de novo* CNVs in affected probands are not depleted of UCEs; rather they are enriched due to overlap with 68 UCEs (Figure 1A, Table S2A; P = 0.024, obs/exp = 1.272). Further, the UCE overlap for Vulto-van Silfhout *de novo* CNVs is significantly different from that of the Vulto-van Silfhout inherited CNVs (Figure 1A, Table S2A; P = 0.002). Therefore, in the ASD patients examined by Vulto-van Silfhout *et al.*^51^, *de novo* CNVs in ASD probands affect an excess of UCEs as compared to inherited CNVs. Finally, we examined the relationship between Vulto-van Silfhout ASD *de novo* CNVs and intergenic, intronic, and exonic UCEs since our previous studies had suggested that intergenic, intronic, and exonic UCEs may differ somewhat in their relationship to CNVs. We found that the enrichment for UCEs is driven by intronic elements, with 36 UCEs overlapping the *de novo* CNVs (Figure 1C, Table S2A; P = 0.010, obs/exp = 1.487).

**Figure 1:**
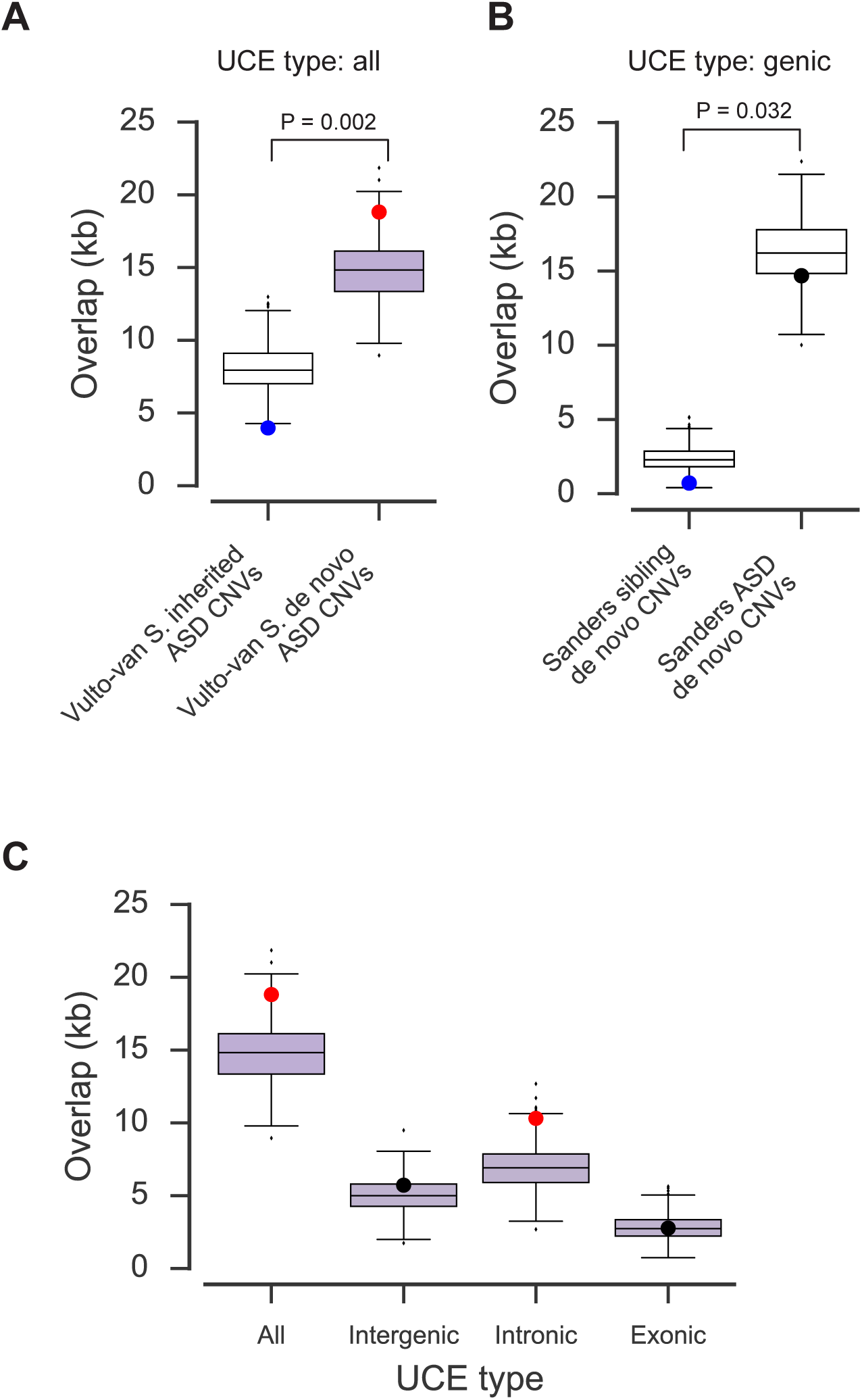
*De novo* CNVs in cases of autism spectrum disorder disrupt UCE dosage. (A): UCEs are depleted from Vulto-van Silfhout ASD inherited CNVs and enriched in Vulto-van Silfhout ASD *de novo* CNVs. Vulto-van Silfhout ASD *de novo* CNVs are represented in lilac throughout the figures. (B): UCEs in genes are depleted from Sanders sibling *de novo* CNVs, and not depleted from Sanders ASD *de novo* CNVs. A X^2^ test for the observed and mean expected numbers of UCE overlaps in both conditions shows a significant difference. (C): Enrichment of UCEs in Vulto-van Silfhout ASD *de novo* CNVs is driven by enrichment of intronic UCEs; intergenic and exonic UCEs were not significantly enriched. All panels: Large dots indicate observed overlap of UCEs, red for enriched, blue for depleted, and black for neither. Boxes indicate median and interquartile range for overlap with UCE-matched random regions, whiskers indicate 1.5 × interquartile range, with all other points plotted as outliers. Brackets: A X^2^ test for the observed and mean expected numbers of UCE overlaps in both conditions shows a significant difference.

We next turned to data from Sanders *et al.*^74^, which presents *de novo* CNV data for autism simplex cases (where only one family member is affected) and their unaffected siblings. We obtained 321 ‘Sanders ASD *de novo* CNVs’ from ASD cases (see Figure S1 for details of filtering steps), which collapsed into 191 distinct non-overlapping CNV regions that, together, covered 8.49% of the genome (Table S1). We likewise retained 86 CNVs from siblings of ASD cases (Figure S1), which collapsed into 79 non-overlapping CNV regions covering 1.15% of the genome, and these CNVs comprise our ‘Sanders sibling *de novo* CNV’ dataset (Table S1).

We found that Sanders ASD *de novo* CNVs and Sanders sibling *de novo* CNVs both fail to show significant depletion of UCEs and thus deviate from the profile that is generally obtained for CNVs representing healthy individuals. Specifically, Sanders ASD *de novo* CNVs overlap 78 UCEs (Table S2A; P = 0.278, obs/exp = 0.938), and Sanders sibling *de novo* CNVs overlap 7 UCEs (Table S2A; P = 0.056, obs/exp = 0.538), with the difference between their degrees of overlap being insignificant (Table S2A; P = 0.123). The result for Sanders sibling *de novo* CNVs presents an intriguing picture. On the one hand, they do not show significant depletion of UCEs and thus resemble the Sanders ASD *de novo* CNVs. On the other hand, their obs/exp ratio of 0.538 approaches the ratio of 0.395 we observed in our previous analysis of *de novo* CNVs representing unaffected individuals^35^.

To further investigate differences between the Sanders sibling and ASD *de novo* CNVs, we next examined intergenic, intronic and exonic UCEs separately. With respect to intergenic UCEs, neither Sanders sibling *de novo* CNVs nor Sanders ASD *de novo* CNVs show depletion (Table S2A; Proportion = 0.354, obs/exp = 1.197, P = 0.425, obs/exp = 1.039 for Sanders sibling and ASD *de novo* CNVs, respectively). This may suggest that the intergenic UCE profiles of both sets of CNVs are not commensurate with that of unaffected individuals. For intronic and exonic UCEs, respectively, Sanders sibling *de novo* CNVs showed low obs/exp values of 0.303 and 0.298 (Table S2A; P = 0.047 for intronic UCEs, proportion = 0.096 for exonic UCEs), while the analogous values for Sanders ASD *de novo* CNVs were higher at 0.835 and 1.054, respectively (Table S2A; P = 0.137 and P = 0.402). Similarly, Sanders sibling *de novo* CNVs are depleted for all genic UCEs, that is, intronic and exonic UCEs, combined (Figure 1B, Table S2A; P = 0.019, obs/exp = 0.302). In contrast, not only do Sanders ASD *de novo* CNVs fail to show depletion of genic UCEs (Figure 1B, Table S2A, P = 0.212, obs/exp = 0. 900) they overlap genic UCEs significantly more than do Sanders sibling *de novo* CNVs (Figure 1B, Table S2A; P = 0.032). This suggests that, for UCEs in genes, *de novo* CNVs in ASD probands show a greater propensity to disrupt UCE copy number than do *de novo* CNVs in unaffected siblings.

In sum, we have found that disruption of UCE copy number by *de novo* CNVs is elevated in two cohorts of individuals with ASD. In one cohort, *de novo* CNVs in ASD probands disrupt UCEs more than do inherited CNVs in the same individuals while, in a second cohort, *de novo* CNVs in ASD probands disrupt UCEs in genes more than do *de novo* CNVs in their unaffected siblings.

### UCEs are strongly enriched in proximity to the breakpoints of balanced rearrangements in subjects with neurodevelopmental phenotypes

Given the importance of balanced rearrangements in the genetic etiology of NDDs,^55, 56, 75–77^ we asked whether the connection between UCEs and NDD-associated genomic rearrangements is limited to CNVs or whether it extends to copy number neutral, or ‘balanced’, rearrangements. Defining balanced rearrangements as those that did not delete or duplicate more than 1 kb of sequence around each breakpoint, we drew our first set of balanced rearrangements drew from five publications that provided the information on breakpoints and phenotypes that is required for our filtration steps: Granot-Hershkovitz *et al.*^53^, Kloosterman *et al.*^54^, Chiang *et al.*^55^, Talkowski *et al.*^56^, and Nazaryan *et al.*^51^ (see Methods, Table S1, and Figure S3 for filtering steps and a list of breakpoints with their respective publication sources).

As expected, no UCEs were positioned directly over any of the breakpoints (likelihood of overlap = 0.012; Methods). We therefore asked whether UCEs are enriched in the vicinity of the breakpoints by querying the 100kb genomic regions flanking either side of the breakpoints. A flank size of 100kb was chosen because it is the smallest flank size that that produces a normal distribution of expected overlaps and can thus provide a P-value for depletion or enrichment of UCEs. When flanks surrounding different breakpoints overlapped, we merged them into contiguous regions. The resulting dataset, called ‘NDD breakpoints with 100kb flanks set 1’ (Table S1, Figure S3) and encompassing 76 genomic regions and 0.51% of the genome, overlaps 22 UCEs and shows strong enrichment for UCEs (Figure 2A, Table S2B; P = 2.22×10^−16^, obs/exp = 4.795).

**Figure 2:**
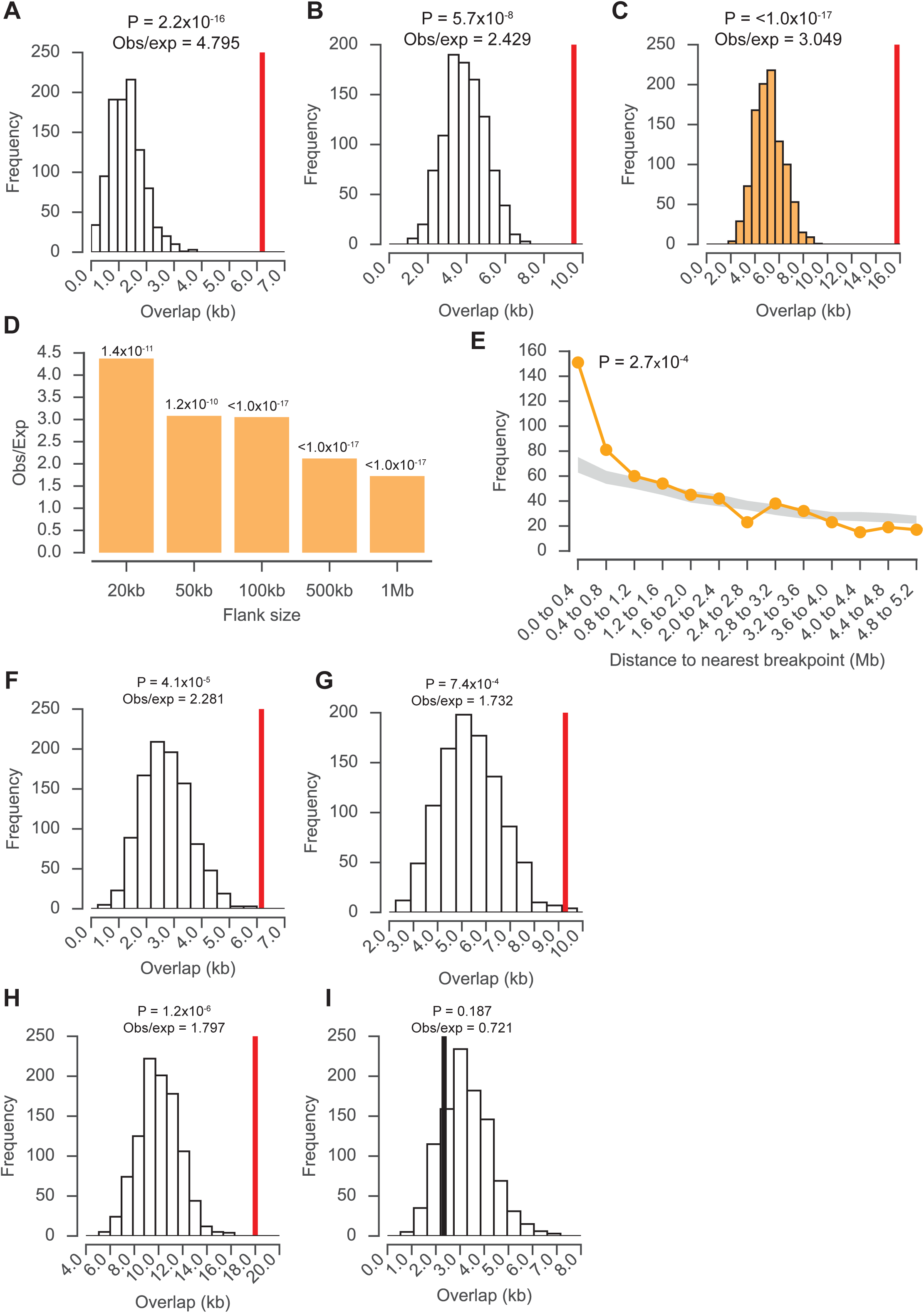
UCEs are enriched in proximity to NDD breakpoints. (A-C): UCEs are enriched in the regions extending 100kb on either side of NDD breakpoints set 1 (A), independent NDD breakpoints set 2 (B) and the combination of the two sets (C). Breakpoints shown in (C) are referred to as ‘Pooled NDD breakpoints’, and analyses using this dataset are in orange throughout. Histogram bars show distribution of expected overlaps, red bars show observed overlap between UCEs and breakpoint flanking regions. (D): UCEs are enriched around pooled NDD breakpoints when five different flank sizes are chosen. P values for the analysis are displayed above the bars. (E): UCEs (orange) are closer to breakpoints than random regions matched to UCEs (grey: inter-quartile range for 1,000 iterations of random regions matched to UCEs). P value is for Anderson-Darling test of distribution of distances from UCEs to breakpoints compared with the distribution of random regions to breakpoints, with a maximum distance of 5.2Mb. (F-H): UCEs were enriched in the regions extending 500kb on either side of breakpoints from set 2, whether the subject is designated as having a ‘pathogenic’ variant (F),‘likely pathogenic’ variant (G), or ‘variant of unknown significance’ (H). (I): UCEs were not enriched in the regions extending 500kb on either side of NDD breakpoints described by Redin *et al.* ^60^ that came from subjects with phenotypes unrelated to neurodevelopment.

We next gathered a set of completely independent breakpoints, again in subjects with NDD and related phenotypes, from Redin *et al.*^60^ (Methods, Figure S3). Again, as no direct overlaps between these breakpoints and UCEs were observed (likelihood of overlap = 0.034; Methods), 100kb flanks were added on either side of each breakpoint. The resulting dataset, called ‘NDD breakpoints with 100kb flanks set 2’ (Table S1, Figure S3) and comprising 224 regions and 1.52% of the genome, is also enriched in UCEs, overlapping 36 UCEs (Figure 2B, Table S2B; P=5.7×10^−8^, obs/exp = 2.429). Combining NDD breakpoint set 1 and 2 into ‘Pooled NDD breakpoints’ (Table S1, Figure S3) produced a dataset of 296 regions and covering 2.01% of the genome that is also significantly enriched for UCEs, overlapping 58 elements (Figure 2C, Table S2B; P <1.0×10^−17^, obs/exp = 3.049).

Because the size of the flanks placed around each breakpoint might affect the degree and significance of UCE enrichment, the Pooled NDD breakpoint dataset was analyzed with four additional flank sizes, two less than and two greater than the original 100kb to give the following series of flank sizes: 20kb, 50kb, 100kb, 500kb, and 1Mb (Table S1, Figure S3). Note that the larger number of breakpoints in the Pooled NDD breakpoints dataset produced normally distributed overlaps even with smaller flanks. With respect to the number of regions covered, the resulting datasets ranged from 316 regions for 20kb flanks to 219 for 1Mb flanks, the larger flanks giving rise to a smaller number of regions because they are more likely to overlap and thus be combined into contiguous regions. With respect to the percentage of genome covered, the datasets ranged from 0.42% for 20kb flanks to 16.59% for 1Mb flanks. In all cases, the datasets were significantly enriched for UCEs (Figure 2D, Table S2B; 1.0× 10^−17^ ≤ P ≤ 1.2× 10^−10^, 1.719 ≤ obs/exp ≤ 4.367). Interestingly, the highest obs/exp ratio was observed for the smallest flank size (Figure 2D; obs/exp = 4.367 for 20kb flanks), and the obs/exp ratios decreased as flank size increased, indicating that the enrichment of UCEs near Pooled NDD breakpoints is most extreme when regions closest to the breakpoints themselves are examined.

Our observations of enrichment suggest that UCEs and Pooled NDD breakpoints tend to be more closely positioned than would be expected by chance. To explore this further, we first calculated the distances from UCEs to the nearest Pooled NDD breakpoint. We then obtained 1,000 sets of control genomic regions, matched to UCEs in terms of number, length, and genomic region of interest (intergenic, intronic, or exonic) using our analysis pipeline described previously. Next, for each of the 1,000 sets, we calculated the distances from the control regions to the nearest breakpoint to produce a distribution of expected distance measurements. We then compared the distributions of observed (UCE-to-breakpoint) distances to the expected (control region-to-breakpoint distances), using an Anderson-Darling test (Methods, “Distances from breakpoints to UCEs”), and found the distributions to be significantly different (Figure 2E; P = 2.7× 10^−4^), reflecting that, at smaller distances, UCEs are more prevalent than are control regions (Figure 2E). These findings argue that UCEs are located significantly closer to Pooled NDD breakpoints than are randomly placed control regions (Figure 2E).

As the subjects described by Redin *et al.*^60^ were classified into three categories based on the predicted pathogenicity of their genomic rearrangements, we asked whether balanced rearrangement breakpoints are differentially positioned with respect to UCEs depending on the classification of the individual. The three categories are: 1) ‘pathogenic’, having strong evidence that the rearrangement is involved in NDD etiology; 2) ‘likely pathogenic’, having somewhat weaker evidence that the rearrangement is involved in NDD etiology; and 3) variants of unknown significance (‘VUS’). For each of the three categories, we added 500kb flanks on either side of all breakpoints; flank sizes of 50kb and 100kb were unable to produce normally distributed expected overlaps due to the small number of breakpoints in each category. The resulting ‘pathogenic’, ‘likely pathogenic, and ‘VUS’ datasets respectively contained 32, 62, and 120 regions and covered 1.04%, 2.11%, and 4.07% of the genome (Table S1; Figure S3). Regions surrounding breakpoints in all three categories are enriched with UCEs, with the pathogenic regions overlapping 25 UCEs (Figure 2F, Table S2B; P = 4.1 ×10^−5^, obs/exp = 2.281), likely pathogenic affecting 33 UCEs (Figure 2G, Table S2B; P = 7.3×10^−4^, obs/exp = 1.732), and VUS encompassing 64 UCEs (Figure 2H, Table S2B; P = 1.2× 10^−6^, obs/exp = 1.797). That even variants of unknown significance display an association with UCE position suggests that proximity to UCEs could be investigated as a novel way to classify and understand these rearrangements.

Finally, we examined the breakpoints of rearrangements from Redin *et al.*^60^ from subjects that did not present with phenotypes related to neurodevelopment, but displayed, instead, other congenital or developmental phenotypes (Methods). Calling this dataset ‘non-NDD breakpoints’, we added 500kb flanks on either side of each breakpoint, as 50kb and 100kb flanks did not produce a normally distributed set of expected overlaps (Figure S3, Table S1). Interestingly, and unlike all datasets representing patients with NDD and related phenotypes, this dataset containing 38 regions and covering 1.25% of the genome is not enriched for UCEs (Figure 2I, Table S2B; P = 0.187, obs/exp = 0.721). That not all copy number neutral breakpoints have elevated numbers of UCEs in their vicinity suggests that local enrichment for UCEs is not a universal feature of breakpoints, and that the enrichment of UCEs surrounding the Pooled NDD breakpoints may be a particular feature of subjects with NDD and related neurodevelopmental phenotypes. In sum, these studies of balanced rearrangements argue that association between UCE position and structural variants is not limited to those that disrupt UCE dosage.

### Gene sets related to neurodevelopment do not explain the excess of UCEs affected by Vulto-van Silfhout ASD *de novo* CNVs or the enrichment of UCEs near breakpoints of balanced rearrangements

We also considered the possibility that the enrichment for UCEs in our Vulto-van Silfhout ASD *de novo* CNV and Pooled NDD breakpoint datasets is driven by UCEs within or near specific subsets of genes. Five groups of genes were tested: 1) ‘LoF genes’, which are defined as those with two or more *de novo* loss-of-function mutations discovered across a cohort of NDD patients and considered by Redin *et al.*^60^ as strong candidates for NDD causation or involvement; 2) ‘Constrained genes’, whose degree of loss-of-function variation in healthy individuals is less than would be predicted by models of mutation rates and thus are considered strong candidates for disease association^61^; 3) ‘Embryonic brain genes’, which are expressed specifically in the embryonic human brain and thus are considered a proxy for genes involved in neurodevelopment^62^; 4) ‘SZ genes’, which are associated with schizophrenia through genome-wide association studies^64^; 5) ‘T2D genes’, which are associated with type 2 diabetes through genome wide association studies^63^. The LoF and Embryonic brain genes were chosen because they are directly connected to processes and pathways relevant to neurodevelopment, and the Constrained and SZ genes were chosen because they have been previously documented to overlap with genes involved in NDD^56, 61, 70^. The T2D genes were chosen as a control for our studies, as they represent a disease that is not thought to involve genomic regions relevant to neurodevelopment. As described below, our studies involved three steps.

Our first step of analysis assessed the frequency with which UCEs are found within the five gene sets, considering intronic and exonic portions of the genome only, using our previously described pipeline (Methods, ‘Depletion and enrichment analysis with gene sets excluded’). We first found that LoF, Constrained, and Embryonic brain gene sets are significantly enriched for UCEs (Figure 3A, Table S2C; P < 1.0× 10^−17^, obs/exp = 4.377 for LoF genes, P < 1.0× 10^−17^, obs/exp = 1.910 for Constrained genes, and P = 3.7× 10^−8^, obs/exp = 1.704 for Embryonic brain genes). For SZ genes, we observed nominal enrichment (Figure 3A, Table S2C; P = 0.025, obs/exp = 1.419). T2D genes do not show significant enrichment, albeit with an obs/exp ratio above one (Figure 3A, Table S2C; P = 0.156, obs/exp = 1.467).

**Figure 3:**
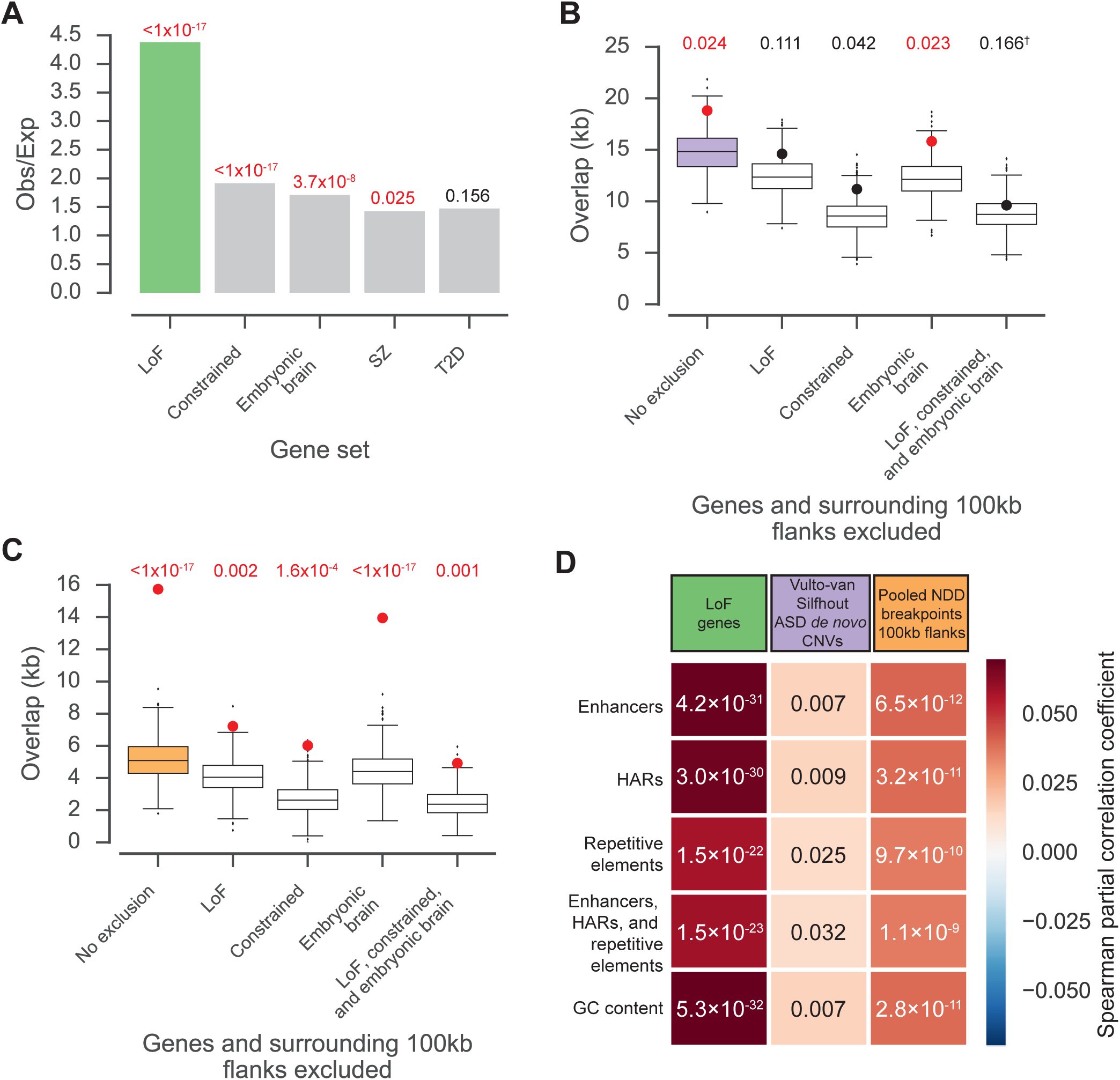
Gene sets related to neurodevelopment do not explain the excess of UCEs affected by Vulto-van Silfhout ASD *de novo* CNVs or the enrichment of UCEs near Pooled NDD breakpoints. (A) UCEs are enriched in genes with 2 or more loss-of-function mutations in ASD subjects (LoF, shown in green in panels A and D), genes with high loss-of-function intolerance scores (constrained), and genes specifically expressed in embryonic brain. UCEs were nominally enriched in genes associated with schizophrenia (SZ) and not enriched in genes associated with type 2 diabetes (T2D). Bar height shows obs/exp ratio. P values are displayed above the bars. Red indicates significant enrichment. (B) UCEs are enriched in Vulto-van Silfhout ASD *de novo* CNVs (lilac box). When regions corresponding to the three most enriched gene sets from panel (A) plus 100kb flanks upstream and downstream of the genes are excluded from the analysis, UCE enrichment is lost in the cases of LoF and constrained genes. For embryonic brain genes, enrichment remains. When all three gene sets are combined, a normal distribution of expected overlaps was not produced, but the proportion of expected overlaps that was greater than or equal to the observed overlap is not consistent with enrichment († symbol). (C) UCEs are enriched in Pooled NDD breakpoint 100kb flanks (orange box). When regions corresponding to the three most enriched gene sets from panel A plus 100kb flanks upstream and downstream of the genes are excluded from analysis, UCE enrichment remains significant. Panels B and C: Large dots indicate observed overlap of UCEs, red indicates significant enrichment, black indicates lack of significant enrichment. Boxes indicate median and interquartile range for overlap with UCE-matched random control regions. Whiskers indicate 1.5x interquartile range, with all other points plotted as outliers. (D) Partial correlation analysis shows that positive correlation between the density of UCEs and LoF genes, and between the density of UCEs and Pooled NDD breakpoints with 100kb flanks is robust to co-correlation with enhancers, human accelerated regions (HARs), repetitive elements, these three combined, and GC content. For Vulto-van Silfhout ASD *de novo* CNVs, correlation with UCEs remained significant when controlling for co-correlation with enhancers, HARs, and GC percentage, but significance was lost when controlling for repetitive elements and for combined enhancers, HARs, and repetitive elements. Heatmap color represents shows spearman partial correlation coefficients; P values are noted within the cells; genome divided into 100kb bins; α = 0.01 considering Bonferonni correction for 5 tests. All panels: green -LoF genes; lilac -Vulto-van Silfhout ASD *de novo* CNVs; orange -Pooled NDD breakpoints.

Because the LoF, Constrained, and Embryonic brain genes display the strongest enrichment for UCEs, we reasoned that they are the most likely to drive enrichment in Vulto-van Silfhout ASD *de novo* CNVs. Here, the genomic regions occupied by, and extending 100kb on either side of, LoF, Constrained, and Embryonic brain gene sets were excluded, both in turn and all at once, from our standard analysis pipeline (Methods ‘Depletion and enrichment analysis’). If the gene sets were responsible for enrichment, then exclusion of these genes and their flanking regions from our analyses would result in loss of enrichment. If such a result were obtained, our studies would further reveal whether the Vulto-van Silfhout ASD *de novo* CNVs would, under these circumstances, resemble that of unaffected individuals and be depleted of UCEs.

Our studies showed that excluding LoF genes and their 100kb flanks caused UCE enrichment to be lost from Vulto-van Silfhout ASD *de novo* CNVs (Figure 3B, Table S2D; P = 0.111, obs/exp = 1.174). A similar outcome was obtained for Constrained genes. (Figure 3B, Table S2D; P = 0.042, obs/exp = 1.306). For Embryonic brain genes and their 100kb flanks, enrichment remained (Figure 3B, Table S2D; P = 0.023, obs/exp = 1. 294). When we removed all three gene sets at once, including 100kb on either side of all genes, our expected overlaps were not normally distributed, and so we did not calculate a P-value using a Z-test. Instead, we found that the proportion of expected overlaps that equaled or exceeded the observed overlap is inconsistent with enrichment (Figure 3B, Table S2D; Proportion = 0.166, obs/exp = 1.191), as is the result of a χ^2^ test to discern whether the number of UCEs overlapped differs from expectation calculated from the genome coverage of the CNVs (Table S2D; P = 0.676). Enrichment was also lost when we excluded the gene sets, either alone or in combination, in all cases without excluding the regions 100kb on either side of the genes (Table S2D). Taken together, these outcomes indicate that the enrichment of UCEs in Vulto-van Silfhout ASD *de novo* CNVs is due in large part to UCEs inside and within 100kb of LoF and Constrained genes. Importantly, however, none of these studies caused the profile of CNVs to be depleted of UCEs (obs/exp ≥ 1.095 in all cases), reinforcing our observations that Vulto-van Silfhout ASD *de novo* CNVs are distinct from CNV datasets representing unaffected individuals. As such, the excess of UCEs affected by the Vulto-van Silfhout ASD *de novo* CNVs continues to suggest an importance of these UCEs in neurodevelopment.

Finally, we turned to the Pooled NDD breakpoints with 100kb flanks. Here, we found that enrichment for UCEs remained after exclusion of LoF, Constrained, or Embryonic brain genes, or all three sets combined, together with their 100kb flanks. While these analyses resulted in larger P-values and smaller obs/exp ratios, suggesting that UCEs in and flanking these genes do contribute to enrichment, they nevertheless demonstrate that such UCEs do not, alone, explain enrichment (Figure 3C, Table S2E; P = 0.002, obs/exp = 1.756 for LoF genes, P = 1.6× 10^−4^, obs/exp = 2.231 for Constrained genes, P = <1.0×10^−17^, obs/exp = 3.143 for Embryonic brain genes, and P = 0.001, obs/exp = 2.031 for all three sets combined). Enrichment was also maintained when we excluded just the gene sets, alone or in combination, but did not extend the exclusion 100kb on either side of the genes (Table S2E). In summary, the enrichment of UCEs in the genomic regions around the breakpoints of balanced rearrangements associated with NDD cannot be explained by UCEs residing in, or within 100kb of, genes implicated in NDD.

### Positive correlation between UCEs and both LoF genes and Pooled NDD breakpoints is robust to other genomic features

We next queried the underlying basis for the enrichment of UCEs in Vulto-van Silfhout ASD *de novo* CNVs (Figure 1A and 3B), LoF genes (Figure 3A), and the vicinity of Pooled NDD breakpoints (Figure 2C and 3C). In particular, partial correlation analyses (Methods) were implemented to examine the contributions of four genomic features: enhancers, regions of the genome that have experienced accelerated sequence change, known as human accelerated regions^78^ (HARs; described further in Discussion), repetitive elements, and GC percentage. We included enhancers because they can overlap UCEs^12, 13, 17^ and HARs because they have been associated with NDD^66^. GC content and repetitive elements were included because they can affect the formation of structural variants^79, 80^. Dividing the genome into 100kb bins and considering α = 0.01 due to Bonferroni correction for 5 tests, we found that, for Vulto-van Silfhout ASD *de novo* CNVs, correlation with UCEs remained significant when controlling for co-correlation with enhancers, HARs, and GC percentage, but was lost when controlling for repetitive elements either alone or in combination with enhancers and HARs (Figure 3D). These results suggest that the association between UCEs and Vulto-van Silfhout ASD *de novo* CNVs is robust to the positions of enhancers, HARs, and to GC percentage, but not to repetitive element position, possibly due to the fact that UCEs are anti-correlated with repetitive elements^33^ and CNVs are positively correlated with repetitive elements^79^.

Having observed an impressive enrichment for UCEs in LoF genes, additional analyses were conducted to address the potential contribution of other genomic features in this context, too. The correlation between UCEs and LoF genes remained significant in all cases, including when the enhancers, HARs, and repetitive elements were combined together. Finally, we examined Pooled NDD breakpoints with 100kb flanks, and found that correlation with UCEs was again significant when controlling for co-correlation with enhancers, HARs, repetitive elements, these three features combined, and GC percentage. This finding indicates that the enrichment for UCEs near Pooled NDD breakpoints is not obviously influenced by the other genomic features we examined and, instead, may be an inherent feature of the position of UCEs.

## Discussion

The present study documents an excess of UCEs in regions affected by ASD *de novo* CNVs and an enrichment of UCEs in the vicinity of balanced rearrangement breakpoints in individuals with NDD and related phenotypes. UCEs have long been associated with gene regulatory functions, particularly for developmentally important Genes^12–14^ (reviewed in Baira *et al.*^81^, Harmston *et al.*^32^, and Fabris *et al.*^82^). Indeed, our results point to the importance of considering longer range interactions (over 100kb) and spatial genome organization^83, 84^ when exploring how UCEs may regulate genes in neurodevelopment.

We^33–35^ and others^31, 36^have also speculated on a different and non-mutually exclusive model for UCE conservation, wherein UCEs contribute to genome integrity. Here, cells would assess the integrity of UCE-containing genomic regions, perhaps by physically pairing allelic UCEs such that disruptions of pairing compromise cell viability and even organismal fitness. As such, cellular assessment and response to disruption (CARD) would provide selective advantage to organisms by reducing their burden of deleterious chromosomal changes^33–35^. Indeed, such a mechanism is consistent with, and may help to explain, the enrichment of UCEs in NDD-relevant genomic regions (Figure S4). Specifically, elimination of deleterious mutations would be most advantageous in regions that are functionally important and/or highly vulnerable to mutation, and NDD-associated regions meet both criteria; the functional importance of NDD-associated regions is evidenced by the severity of many NDD phenotypes, while the vulnerability of genes involved in neuronal and brain development and function is exacerbated by their longer than average length^85–87^, subjecting them to increased replication stress and thus chromosome rearrangements (Wilson *et al.* ^88^ and references therein).

Curiously, a recent study has reported that NDD-relevant regions contain an excess of sequences, known as HARs, that had been highly conserved and then, in humans, experienced accelerated rates of change^66^. How might this paradoxical situation arise, wherein a highly conserved sequence, perhaps a UCE, suddenly begins to change at an accelerated rate? We note first that mutations to UCEs can, in fact, acquire SNPs^89–91^, suggesting that evolution can in some cases erode UCEs. Secondly, heterozygosity has been correlated in certain studies with increased mutation rates, with pairing considered as a possible mechanism for the detection of heterozygosity in some instances^92–94^. Thus, in the case that heterozygosity can increase mutation rate and, in addition, considering that the first instance of an altered UCE will almost certainly result in heterozygosity, this initial mutation might be accompanied by accelerated further mutation of the region. In any case, should alteration of a UCE produce an advantageous function, such as a new regulatory activity, selection for the new function may ultimately release that UCE sequence from the its previous constraints, creating a HAR. As this process might be expected to be most prevalent in genomic regions where UCEs are enriched, such as NDD-relevant regions, it could explain the excess of HARs in these same regions.

To conclude, our results demonstrate an association between neurodevelopmental phenotypes and an elevated level of structural variation affecting UCE dosage or genomic position. Interestingly, a positive correlation has been observed between NDD and an elevated risk for cancer (Norwood *et al.*^95^ and references therein), perhaps reflecting that disruption of UCEs can misregulate genes involved in development and oncogenesis. In light of our previous observations that cancer-specific CNVs are enriched in UCEs^35^, it may also be that individuals with NDDs would lack the proposed safeguard of intact and correctly positioned UCEs^33–35^ later in life, and therefore show higher cancer prevalence.

## Note

During preparation of this manuscript, two manuscripts of relevance were released on the bioarchive preprint server. Firstly, Short *et al.* ^96^ report an enrichment of *de novo* single nucleotide variants (SNVs) in conserved, putatively regulatory, noncoding genomic regions in the genomes of probands with developmental disorders. Secondly, Werling *et al.* ^97^ describe an intriguing excess of *de novo* SNVs and small (<50bp) insertion-deletion (indel) variants within conserved regions in ASD cases compared to controls, though the statistical significance of this observation did not survive multiple testing correction.

## Supplementary Data

The supplementary data consists of four figures and three excel tables.

## Acknowledgements

We thank D. Balick, R. Collins, S. Erdin, K. Mattioli, V. Pillalamarri, S. Sunyaev, R. Tarnita, T. Tullius, D. Weghorn, and all members of the Wu laboratory for helpful and insightful discussions. This work was supported by a William Randolph Hearst Foundation Award to R.B.M., an EMBO Long Term Fellowship to J.E., awards to M.E.T. from NIH (MH095867 and GM061354), the Simons Foundation for Autism Research (SFARI #346042), and the March of Dimes, and awards to Ct.W. from NIH (DP1GM106412; R01HD091797) and Harvard Medical School.

The authors declare that no conflicts of interest exist.

We apologize to authors whose work we were unable to cite due to limits on citation number.

## Supplementary information legends

The supplementary information consists of four figures and three excel files.

### Figures

**Figure S1:**
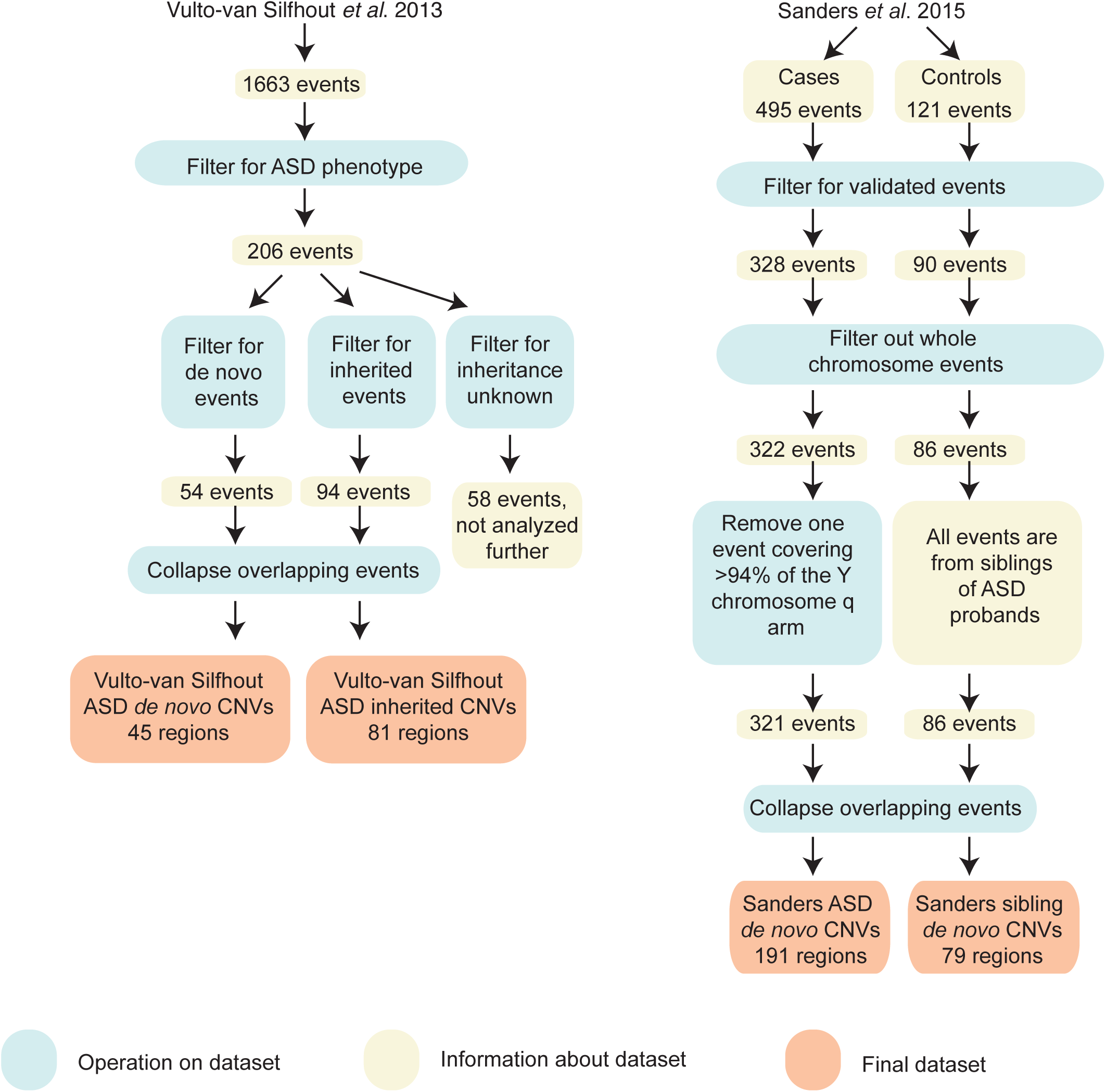
Explanation of methods to obtain datasets. A: Datasets from Vulto-van Silfhout *et al.* B: datasets from Sanders *et al.*

**Figure S2:**
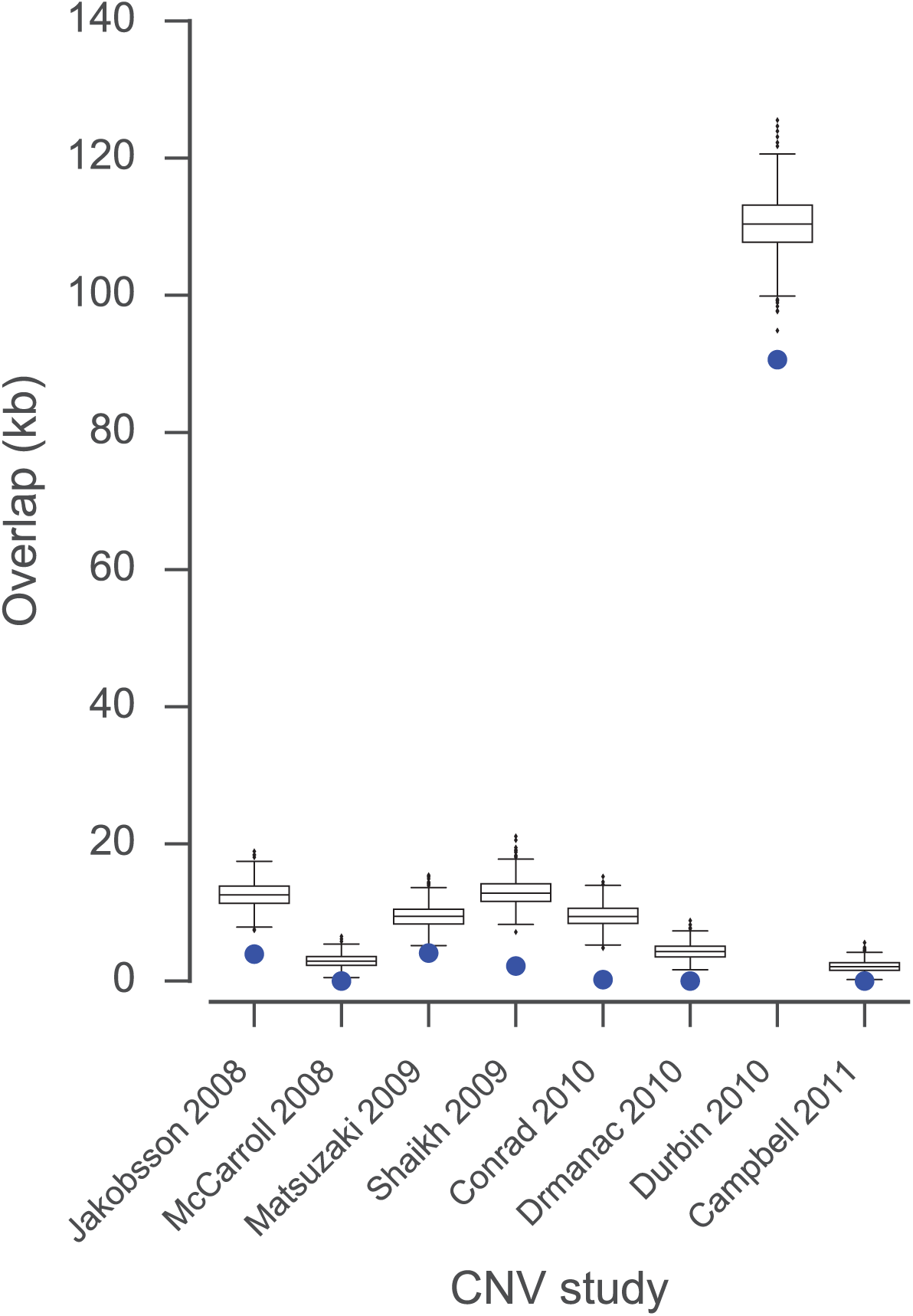
CNVs from healthy individuals are depleted of UCEs. CNV datasets from eight separate studies of healthy individuals were previously analyzed in McCole *et al.* and consistently showed significant depletion for UCEs.

**Figure S3:**
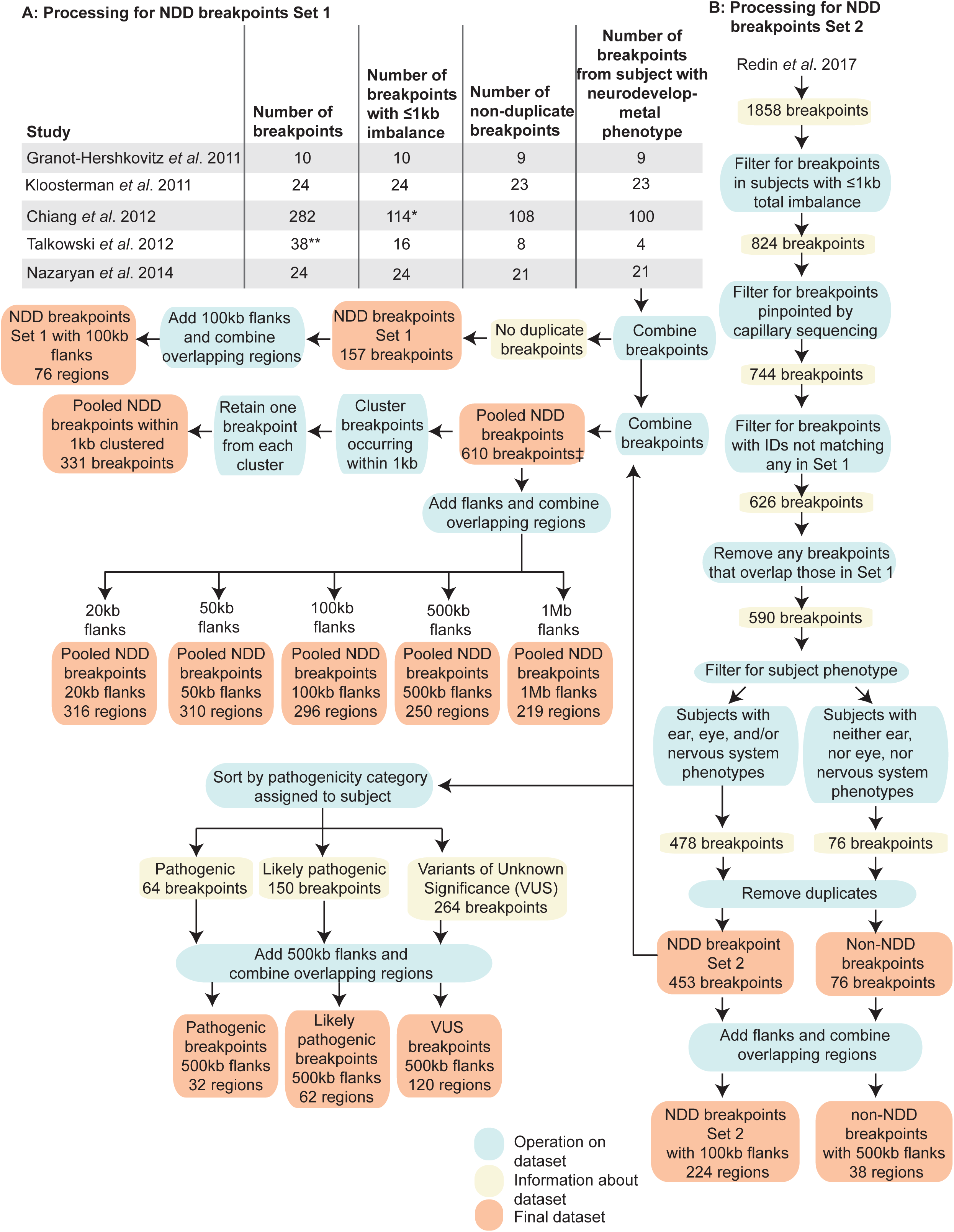
Explanation of methods to obtain breakpoint datasets. A: Processing for NDD breakpoint set 1. *Breakpoints derived from patients where their total genomic imbalance was ≤1kb. **Breakpoints also described in Chiang *et al.* 2012 were removed from the set taken from Talkowski *et al.* 2012. B: Processing for NDD breakpoint set 2.

**Figure S4:**
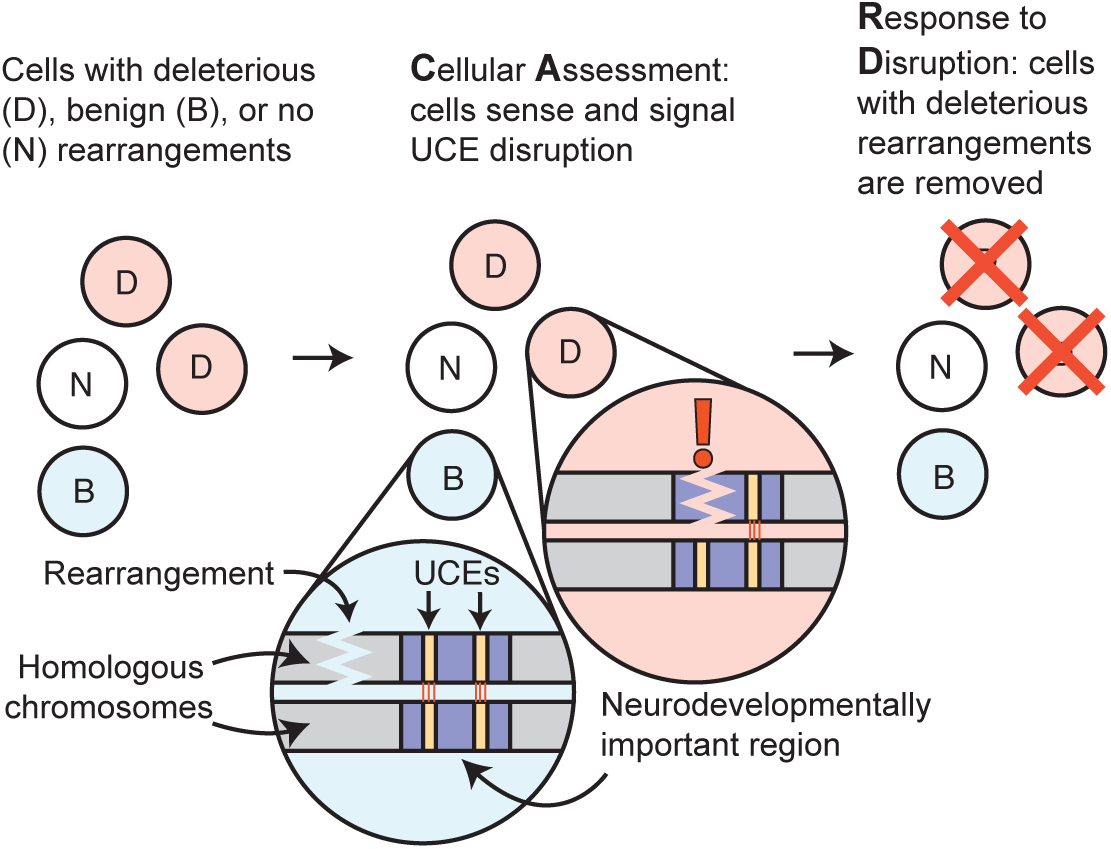
Explanation of the ‘CARD’ mechanism. Cell populations include those with no genomic rearrangements (‘N’, white), with benign rearrangements (‘B’, blue), or with deleterious genomic disruptions (‘D’, pink). Benign rearrangements are positioned where they do not disrupt homologous UCE pairing and comparison (orange lines between yellow UCEs), while deleterious disruptions are those that interfere with UCE pairing and are detected (orange exclamation point). Cells containing deleterious rearrangements are culled.

### Excel files

**Table S1:Datasets.** Tab A: Information on all datasets studied. Subsequent tabs: The coordinates for each dataset listed in (A) are contained in a tab, with the dataset name corresponding to the tab title. Last tab: detailed source information is given for Pooled NDD breakpoints.

**Table S2:Enrichment analysis.** A: CNVs in ASD subjects and siblings B: Balanced rearrangement breakpoints C: Gene sets. D: Vulto-van Silfhout ASD *de novo* CNVs, excluding gene sets from the analysis. E: Pooled NDD breakpoints with 100kb flanks, excluding gene sets from the analysis. All tabs: Proportion: of 1,000 expected overlap iterations, the number of times the expected overlap generated was equal to, or more extreme than, the observed UCE overlap (bp), divided by the total number of iterations, which was always 1,000. P-value: significance of whether the observed overlap (bp) differs from the expected overlaps, as determined by a Z-test. obs/exp: observed overlap (bp) divided by mean of expected overlaps (bp). Outcome: Determined with a two-tailed test (P ≤ 0.025 in each tail for an overall a of 0.05).

**Table S3:List of UCE coordinates together with their overlaps with genes and with all datasets analyzed in this study.**

